# SwrA as global modulator of the two-component system DegS/U in *B. subtilis*

**DOI:** 10.1101/2021.05.07.443137

**Authors:** Francesca Ermoli, Giulia Vitali, Cinzia Calvio

## Abstract

The two-component system DegS/U of *Bacillus subtilis* controls more than one hundred genes involved in several different cellular behaviours. Since the consensus sequence recognized by the response regulator DegU has not been clearly defined yet, mutations in either component have been crucial in the identification of the cellular targets of this regulatory system. Over the years, the *degU32*^Hy^ mutant allele, that was supposed to mimic the activated regulator, has been commonly used to define the impact of this TCS on its regulated genes in domestic strains.

SwrA encodes a small protein essential for swarming motility and for poly-γ-glutamate biosynthesis and is only present in wild strains. Previous work indicated that SwrA is partnering with DegU~P in exerting its role on both phenotypes.

In this work, inserting a *degS200*^Hy^ mutation in *swrA*^+^ and *swrA*^-^ isogenic strains we demonstrate that SwrA modulates the action of DegU~P on two new phenotypes, subtilisin expression and competence for DNA uptake, with a remarkable effect on transformation. These effects cannot not be appreciated with the DegU32^Hy^ mutant as it does not mirror the wild-type DegU protein in its ability to interact with SwrA.

## INTRODUCTION

Two-component systems (TCS) are signal transduction modules common in bacteria and archaea, composed by a sensor histidine kinase and a cognate response regulator. Sensor kinases auto-phosphorylate themselves on a histidine residue in response to specific environmental signals and then transfer the phosphate group to a specific aspartic acid residue of the regulator inducing a structural rearrangement that enables it to modify its DNA binding properties and regulate gene expression. Moreover, sensor kinases can often quench spurious signals by dephosphorylating their cognate regulators (1). The DegS/U TCS, composed by the cytoplasmic DegS kinase and the DegU transcription factor, is involved in the regulation of several important physiological pathways of *Bacillus subtilis,* among which flagella-mediated motility, degradative enzyme synthesis, genetic competence, and sporulation (2). The extremely wide impact of this TCS has been evidenced through several transcriptional profiling experiments (3–5). A particular class of mutations in either DegS or DegU leads to the hyperproduction of several degradative enzymes, including the *aprE*-encoded protease subtilisin, and has therefore been named “Hy” (6–8). Besides promoting the synthesis of several degradative enzymes, the pleiotropic Hy mutations also cause the so called Hy phenotype which includes loss of DNA competence, absence of flagella, sporulation in the presence of glucose and elongated cell morphology (6, 9, 10). A Hy phenotype is also observed when two small proteins, DegQ and DegR, are overexpressed. They are both involved in the DegS/U signalling pathway: DegQ stimulates the transfer of the phosphate moiety from DegS to DegU (5), while DegR stabilizes DegU~P by preventing DegS-mediated DegU~P dephosphorylation (11). The overexpression of DegQ naturally occurs in wild *B. subtilis* strains, thanks to a nucleotide change in the −10 box of its promoter that leads to 10-fold increase in transcription with respect to domestic strains (12, 13); however, the *degQ*^Hy^ mutation present in undomesticated strains only generates a mild phenotype, as these strains, differently from *degS/U*^Hy^ mutants, do not copiously produce γ-PGA (see below).

Among the number of originally isolated Hy mutants (6), subsequent studies have heavily relayed on *degU32*^Hy^, a particular *degU* allele which carries an A-to-T transversion at nucleotide 35 of the *degU* ORF, leading to a His to Leu amino acid change at position 12 (14). In early studies, all DegU32^Hy^ phenotypes matched those obtained with the DegS200^Hy^ mutant, in which the Gly to Glu mutation at position 218 of DegS impairs the phosphatase activity of the sensor kinase, thus leading to the accumulation of DegU~P (15). From the perfectly overlapping phenotypes of the two mutants, DegU32^Hy^ has been since considered as a constitutively active proxy of DegU~P, without any further structural characterization. Only recently such interpretation has been challenged, thanks to the introduction of a new player, SwrA (16).

SwrA is also a small protein, 117 aa, which has been discovered thanks to its fundamental role in swarming motility (17). Its existence remained long undisclosed due to the fact that domestic *B. subtilis* strains only encode a non-functional 13 aa truncated peptide because of the insertion of an extra adenine in a poly-A tract in *swrA* ORF causing a frameshift mutation (17–19). This type of mutations can easily flip back to the wild-type form (wt) and vice versa with the frequency of a phase-variation event (17). It is thus frequent to obtain mixed *swrA*^+/-^ populations even in laboratory strains upon prolonged incubations. When functional, SwrA stimulates flagella production through its activity at the P_fla/che_ promoter, thereby promoting *sigD* transcription which also permits efficient cell septation (16, 19–21). Although initially confined to motility regulation, the role of SwrA was also shown to be essential for the expression of the otherwise-silent *pgs* operon, encoding the enzymes required for the synthesis of the biopolymer poly-γ-glutamic acid (γ-PGA) (22). Interestingly, γ-PGA production not only strictly depends on SwrA but also on a Hy mutation in either DegQ (23) or DegU/S (24). This was the first evidence of a strict connection between SwrA and the DegS/U TCS.

Further genetic evidence of the link between SwrA and the DegS/U was gained by studies on the *fla/che* operon, in which the direct interaction between DegU~P and SwrA was demonstrated genetically and biochemically. Genetically, it was observed that while *degU32*^Hy^ and *degS200*^Hy^ mutants completely suppress P_fla/che_ expression in laboratory strain lacking SwrA, leading to the classically non-flagellated Hy-phenotype (6, 10), the restoration of a functional *swrA* allele leads to hyperflagellation in *degS200*^Hy^ mutants as well as in *degS/U*^wt^ backgrounds, although not in *degUS32*^Hy^ mutants (16). Biochemically, it was shown that, in electro mobility shift assays, the DegU~P-bound *fla/che* promoter is super-shifted in the presence of SwrA while DegU32^Hy^ is not; moreover, DegU32^Hy^ does not require phosphorylation for DNA binding (16). The physical interaction between DegU and SwrA was also evidenced in other studies (25). Ultimately, the impact of SwrA on motility is to remarkably turn the P_fla/che_ repressive effect of DegU~P, naturally produced or induced by a *degS200*^Hy^ mutation, into a transcriptional boost, thus allowing swarming motility (16, 21). This dramatic overturn of DegU~P impact on the motility operon could not be appreciated in domestic strains, because of the absence of SwrA and in wild strains if the *degU32*^Hy^ allele is used as a proxy of DegU~P (26).

Although the above data suggest that SwrA does not interact with DegU32^Hy^, this is not always true. Indeed, γ-PGA production is induced by SwrA in the presence of either *degU32*^Hy^ or *degS200*^Hy^ (24). However, a deep characterization of the differential impacts of the two Hy alleles on Ppgs has yet to be conducted.

In competence, Hy mutants have been shown to have a negative impact on the overall process in laboratory strains. However, *degS/U* null mutants were also shown to be non-competent, suggesting the requirement of this TCS in the pathway (9). The current model is the following: DegU~P has a negative impact on *comS,* while unphosphorylated DegU is required, possibly because it mediates the binding of ComK to its own promoter (27). More recently, the *degQ*^Hy^ allele was shown to negatively affect *comS* and *comK* expression in both domestic and wild strains (28).

In this work we demonstrate that SwrA heavily impacts not only motility and γ-PGA production but also other DegS/U regulated behaviours; SwrA positively modulates DegS/U activity in competence for transformation and reduces *aprE* transcription. Moreover, we characterized the differential influence of *degU32*^Hy^ and *degS200*^Hy^ mutations on *pgs* transcription. Finally, we once more demonstrate that the *degU32*^Hy^ allele encodes a constitutively active mutant protein whose activity dramatically differs from the phosphorylated DegU~P protein. Our results suggest that, as it happened in the past for motility, the use of this mutant may lead to misleading interpretations of the real physiological role of DegS/U TCS in *B. subtilis* physiology.

## RESULTS

### SwrA and motility in undomesticated strains

Our previous work showed that SwrA acts by subverting the impact of DegU~P on the *fla/che* promoter, transforming its action into a positive boost on flagellar gene expression. The functional interaction SwrA-DegU~P only occurs with the wild-type phosphorylated form of the response regulator, while the DegU32^Hy^ mutant protein does not effectively interface with SwrA at this promoter (16). To generalize this effect also to undomesticated strains, either *degU32*^Hy^ or *degS200*^Hy^ mutation was introduced in the transformable *com/*^Q12L^ mutant of the undomesticated NCIB3610 (29). The introduction of the DegU32^Hy^ mutation caused a complete loss of motility, as already shown by Stanley-Wall and collaborators (26). Conversely, the *degS200*^Hy^ derivative of the undomesticated strain proficiently swam and swarmed, paralleling the results obtained in domestic strain (Fig. S1). The only difference with domestic strains is the presence of a well-defined “lump” of γ-PGA that can be observed in the central part of the *degS*^Hy^ plates in Fig. S1. This characteristic is due to the abundant production of the polymer in DegS200^Hy^ mutants, which is much higher than in the *degU32*^Hy^ background. Interestingly, γ-PGA production was never visible in *degS/U*^wt^ undomesticated strains, although they naturally contain the *degQ*^Hy^ mutation. This finding suggests that the *degQ*^Hy^ mutation, which does not impact on the protein structure of DegU, is less effective than the *degS200*^Hy^ mutation in generating DegU~P.

Therefore, we concluded that the powerful overturn of the DegU~P action on motility genes is a general phenomenon occurring not only in laboratory strains, but also in wild, undomesticated strains, even when the phosphorylation of DegU is maximal.

### SwrA and competence for DNA uptake

The voluminous literature data reporting the negative effect of Hy mutations on competence were acquired in domestic *B. subtilis* strains which lack the SwrA protein. To establish whether SwrA has a general role as regulatory factor for DegS/U activity in competence, genetic transformation was analyzed in isogenic mutants differing for the status of the *swrA* allele as well as for the source of DegU~P: either the intact phosphoprotein obtained in the presence of a *degS200*^Hy^ or the mutant DegU32^Hy^ protein.

Transformation efficiency was assessed in PB5370 and PB5249, respectively the *swrA*^-^ and *swrA*^+^ versions of the commonly used JH642 domestic strain (30), which do not contain any selectable marker. The *swrA*^-^ and *swrA*^+^ isogenic strains did not show any significant difference in transformation efficiency (Fig. 1), but for a slightly better performance of the *swrA*^+^ strain, PB5249, which was thus taken as reference strain for determining the efficiency of the others. Both strains were transformed with the Hy mutation in either *degU* or *degS* and the resulting Hy mutants were transformed with a selectable genomic DNA. As shown in Fig. 1, consistently with literature data, in *swrA*^-^ strains transformation efficiency was abolished by both *degU*^Hy^ and *degS*^Hy^ mutations (efficiency 0.7% and 2.7%, respectively). Even in the presence of a functional *swrA* allele the *degU*^Hy^ strain did not substantially modify competence, i.e., the *swrA*^+^ *degU32*^Hy^ strain remained non-transformable (0.7% efficiency). However, the presence of SwrA in the *degS*^Hy^ strain was sufficient to restore competence to 36% of efficiency (Fig. 1).

**Fig. 1.**
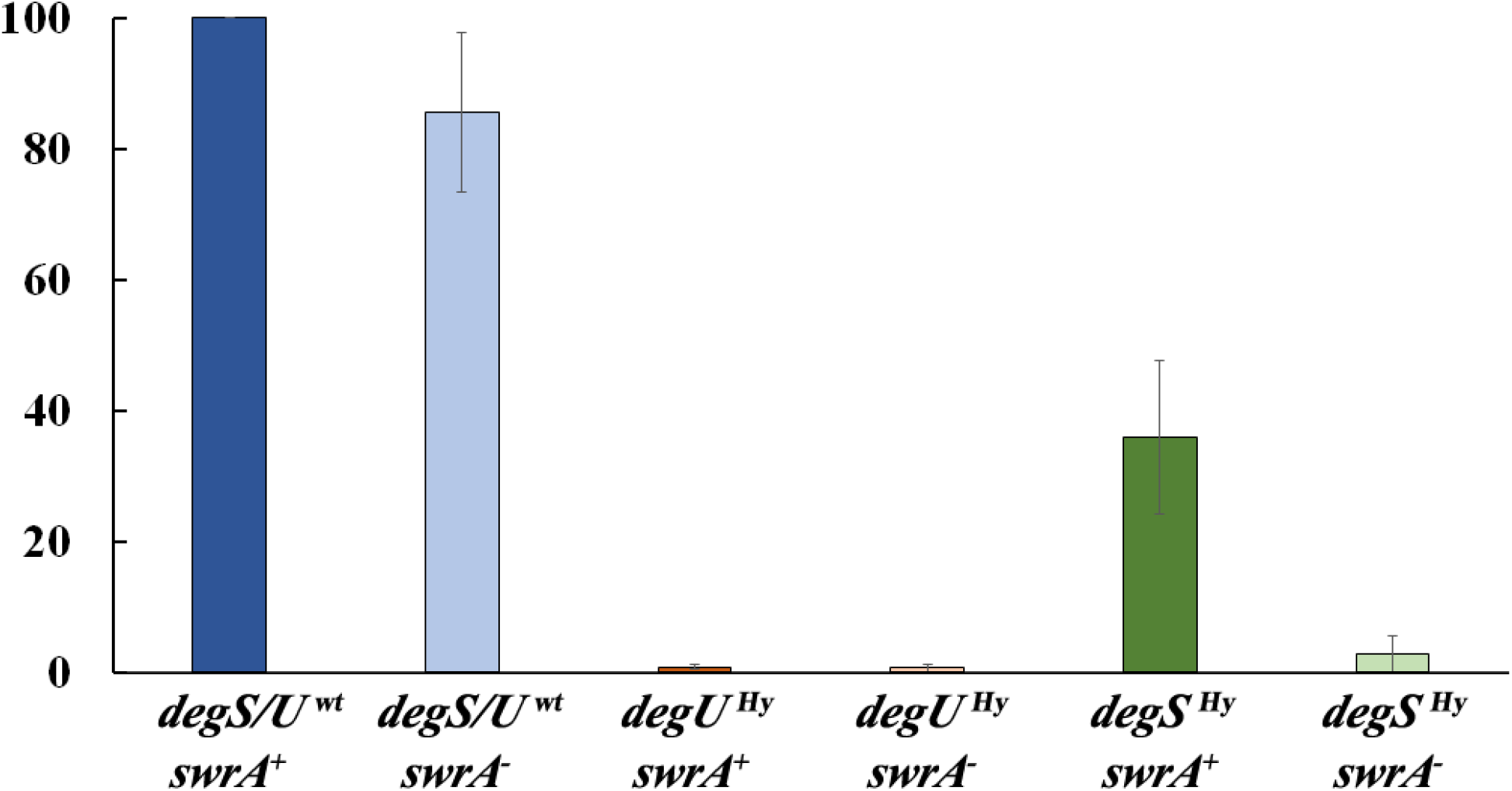
Transformation efficiency. Competence was evaluated in laboratory strains derived from JH642 (*swrA*^-^ *degQ*^wt^ *trpC1 pheA2*), differing for the status of the *degS/U* and *swrA* alleles (listed in Table 1), by counting resistant colonies obtained by transformation with selectable genomic DNA. Each experiment is the average of at least three independent replicates; error bars account for the standard error of the mean. The genotype of each strain is given on the x-axis. In each experiment, 100 % efficiency was assumed for the *swrA*^+^ strain (PB5249).

Taken together, these data indicate that if SwrA is functional, competence is reduced but no longer abolished by phosphorylation of DegU. Thus, SwrA is able to modulate the activity of DegU~P, partially suppressing its negative effect. This positive action is possible only in the presence of a *degS*^Hy^ mutation, i.e., in the presence of a phosphorylated wild-type DegU protein.

### SwrA and subtilisin expression

The restoration of competence in a *degS*^Hy^ strain prompted us to extend our investigation to *aprE* expression, which is known to be induced by the presence of a Hy mutation in either *degS* or *degU* (31). As already described, *swrA*^+^ revertants often arise in the swrA^-^ population upon long incubations and might generate confusing results. To avoid the development of such revertants, a *swrA* null mutant was created together with an isogenic *swrA*^+^ strain. To verify whether SwrA has a role also in subtilisin production, the P_aprE_-GFP reporter, developed by Veening et al. (32), was inserted in *swrA*^+^ and *ΔswrA* JH642-derived strains. Analyses were carried out by imaging flow cytometry, which not only allows to quantify the average single cell fluorescence, but also to dissect the *aprE*-ON and -OFF populations due to heterogeneity in *aprE* expression (32, 33). As the expression of the reporter was not detected in these conditions (data not shown), a *degU32*^Hy^ or *degS200*^Hy^ allele was introduced in each strain. The analyses of the P_aprE_-GFP *degS*^Hy^ or *degU*^Hy^ in both *swrA*^+^ and *ΔswrA* were focused on the transition phase (T_0_), and 5 and 15 h later (T_5_ and T_15_). As shown in Fig. 2A, in DegU^Hy^ strains, the percentage of *aprE*-ON cells did not substantially vary in dependence of the presence of SwrA both at T_0_ and at later time points. Conversely, in DegS^Hy^ strains the percentage of *aprE*-ON cells was highly affected by SwrA. The presence of SwrA led to a substantial decrease in the number of ON cells, particularly at T_0_ (−70%). Moreover, the percentage of ON cells was similar between DegU32^Hy^ or DegS200^Hy^ mutants in *ΔswrA* strains, but it was significantly reduced in the *swrA*^+^ *degS200*^Hy^ background.

**Fig. 2.**
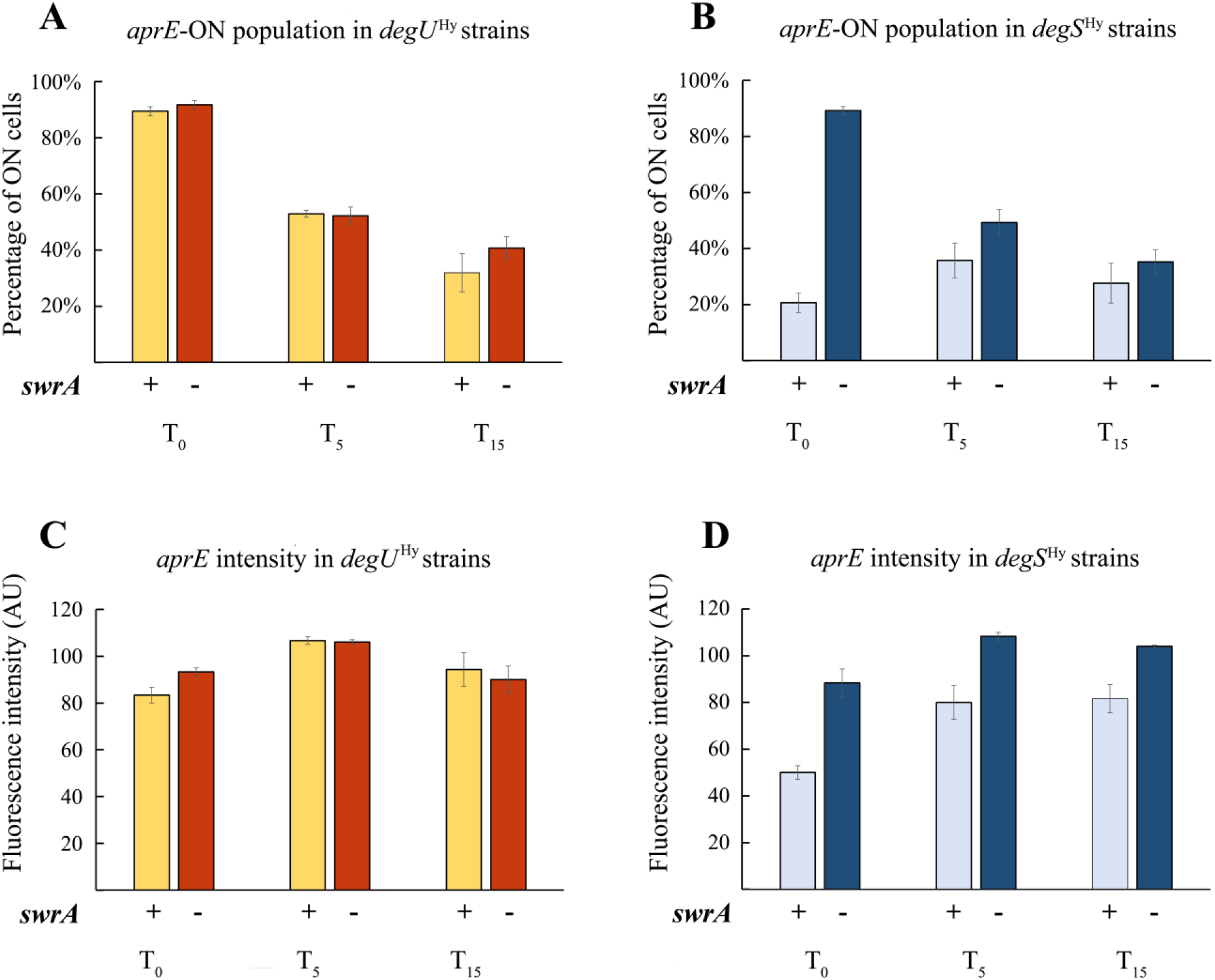
Expression of P_aprE_-GFP. Domestic strains, differing for the status of the *degS/U*^Hy^ alleles and for the presence of a functional *swrA* gene, were analysed by imaging flow cytometry to evaluate the percentage of GFP-ON/OFF cells and the peak of fluorescence intensity. Cultures were sampled at the transition point (T_0_), 5 and 15 hours later (T_5_ and T_15_), as indicated below each graph, where the *swrA* status is also indicated. A, C: data collected for the *degU*32^Hy^ strains; B, D: data collected for *degS*200^Hy^ strains. The upper panels, A and B, represent the percentage of cells expressing the reporter gene (ON-population). The lower panels, C and D, show the peak of intensity of the ON-population. Values represent the average of at least three independent replicates; error bars account for the standard error of the mean.

Also, the expression level of P_aprE_, i.e., the single cell fluorescent intensity, did not vary in the presence or absence of SwrA in DegU^Hy^ strains; however, in DegS^Hy^ mutants the presence of SwrA significantly decreased fluorescence intensity (Fig. 2B). Also in this case, there are no appreciable differences in GFP levels among *ΔswrA* strains, while in the *swrA*^+^ background the *degS200*^Hy^ allele is not as efficient as *degU*^Hy^ in driving *aprE* expression. A gallery of images of *aprE* ON and OFF cells acquired during flow cytometry are provided in Fig. S2.

These data allow to conclude that SwrA modulates the activity of DegU~P also at the *aprE* promoter. SwrA reduces the efficacy of DegU~P on subtilisin expression. Analogously to what observed in competence, the SwrA-mediated effect only occurs in the presence of a *degS*^Hy^ mutation, i.e., in the presence of a wild-type phosphorylated DegU protein.

### DegS^Hy^ and DegU^Hy^ mutants in *pgs* expression

The activation of the *pgs* operon expression is known to depend on the co-presence of at least a *degS*/U^Hy^ allele and SwrA. However, so far, most of the data have been collected using DegU^Hy^ mutants, while scant information is given on γ-PGA production in DegS^Hy^ strains (24). To fill this gap, a P_pgs_-sfGFP reporter construct was devised and inserted *in locus* in the *swrA*^+^ laboratory strain. Since no fluorescence was detected in this strain (data not shown), it was further transformed with either *degS*^Hy^ or *degU*^Hy^ alleles. The Hy strains were grown under vigorous shaking in a glutamate-rich medium that supports γ-PGA production, with periodic sampling over a 48-h prolonged incubation. At relevant time-points, P_pgs_ expression was quantified by imaging flow cytometry. In the DegU^Hy^ mutant, P_pgs_ appeared to be homogeneously active from the beginning of the analysis (2-h post inoculum, data not shown), with intensity reaching a peak at T_-2_ (8-h post inoculum). This early peak of maximal intensity was followed by a monomodal decline over the next 40 h (Fig. 3A), with the majority of the population already OFF after T_18_. Conversely, in the DegS^Hy^ strain, P_pgs_ activation showed a 2-h delay: cells started displaying fluorescence at T_0_ (10-h post inoculum), with a gradual increase over time. Intensity reached a peak at T_14_ which was followed by a slower decline of the GFP signal, which remained however appreciable, in most of the cell population, up to the end of the experiment (T_38_) (Fig. 3B).

**Fig. 3.**
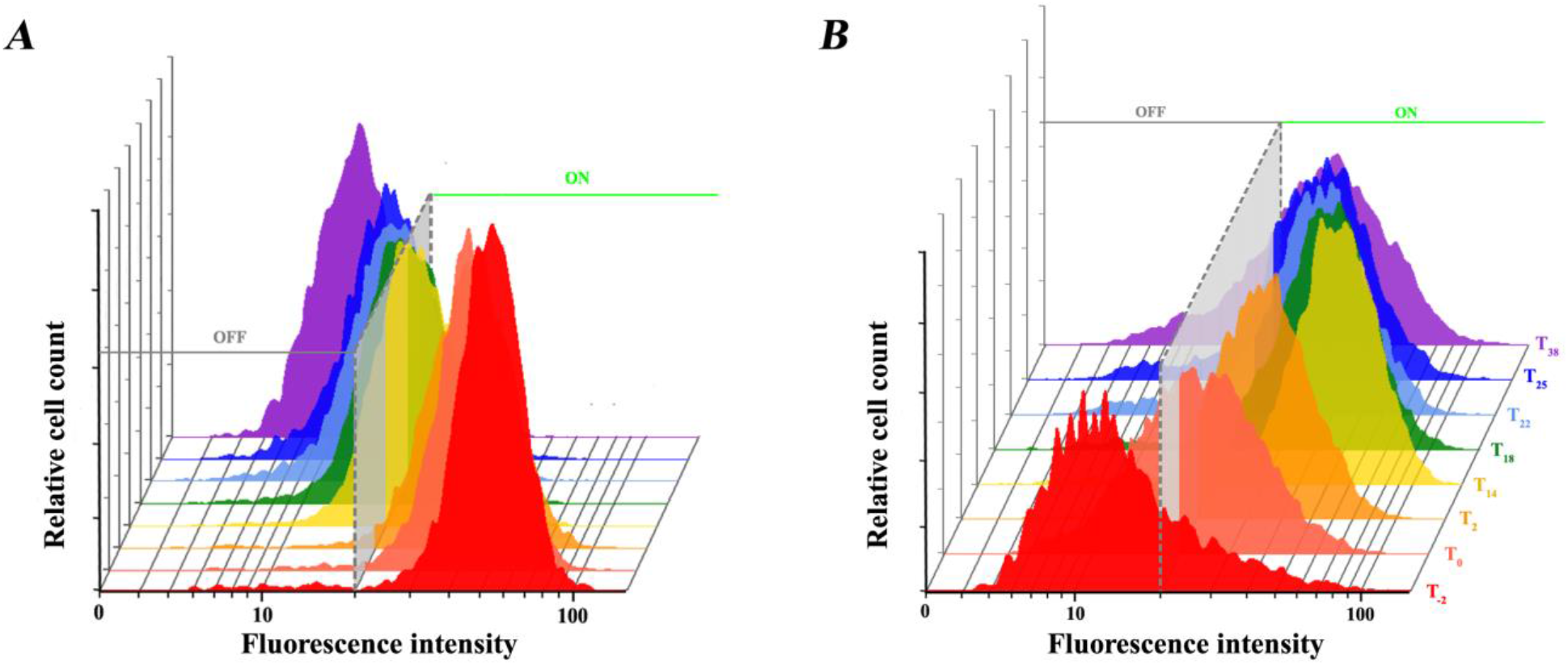
Expression profile of P_pgs_-GFP. Strains (A) *swrA*^+^ *degU*^Hy^ and (B) *swrA*^+^*degS*^Hy^ were grown in E-medium and cells were collected for imaging flow cytometry at different time points. The colour of the plots refers to the collection time (in h relative to the transition point), which is indicated in each graph. A dashed line marks the ON threshold. The intensity values, represented in logarithmic scale on the x-axis, refer to the median pixel intensity of each single event.

From these data it emerges that the impact of SwrA on *pgs* transcription is dramatic for both Hy mutants: no transcription is observed in swrA^-^ backgrounds (22–24), and data not shown). However, there is a remarkable difference in the expression profile using the two partner proteins; upon interaction with SwrA, the constitutively active mutant protein DegU32^Hy^ immediately exerts its pressure on the *pgs* promoter but the effect is rapidly relieved. In the *degS*^Hy^ strain, a delay in *pgs* activation is observed, most likely due to the requirement of the physiological trigger of the signalling pathway. However, once activated, the SwrA-DegU~P stimulus on P_pgs_ is sustained up to 24 h (T_14_), although the intensity of the fluorescent signal is reduced with respect to what observed in cells containing DegU^Hy^. These data are in line with our experimental evidence that DegS^Hy^ strains produce a much higher amount of γ-PGA (data not shown) and indicate that, although an interaction between SwrA and DegU^Hy^ occurs, the effect on transcription is considerably different from that obtained when SwrA interacts with DegU~P.

## DISCUSSION

This work extends the array of DegS/U regulated phenotypes in which SwrA plays a pivotal role. The data have been summarized in Table 2. Considering the phenotypes thus far analyzed, SwrA emerges as key modulator of DegS/U on all the promoters tested so far, P_aprE_, P_pgs_ (Figs. 2 & 3) and P_fla/che_, (16) (Fig. 4). Notably, SwrA also mitigates the negative effect of DegU~P on genetic competence (Fig. 1) and makes *degS*^Hy^ *swrA*^+^ strains easily transformable.

**Fig. 4.**
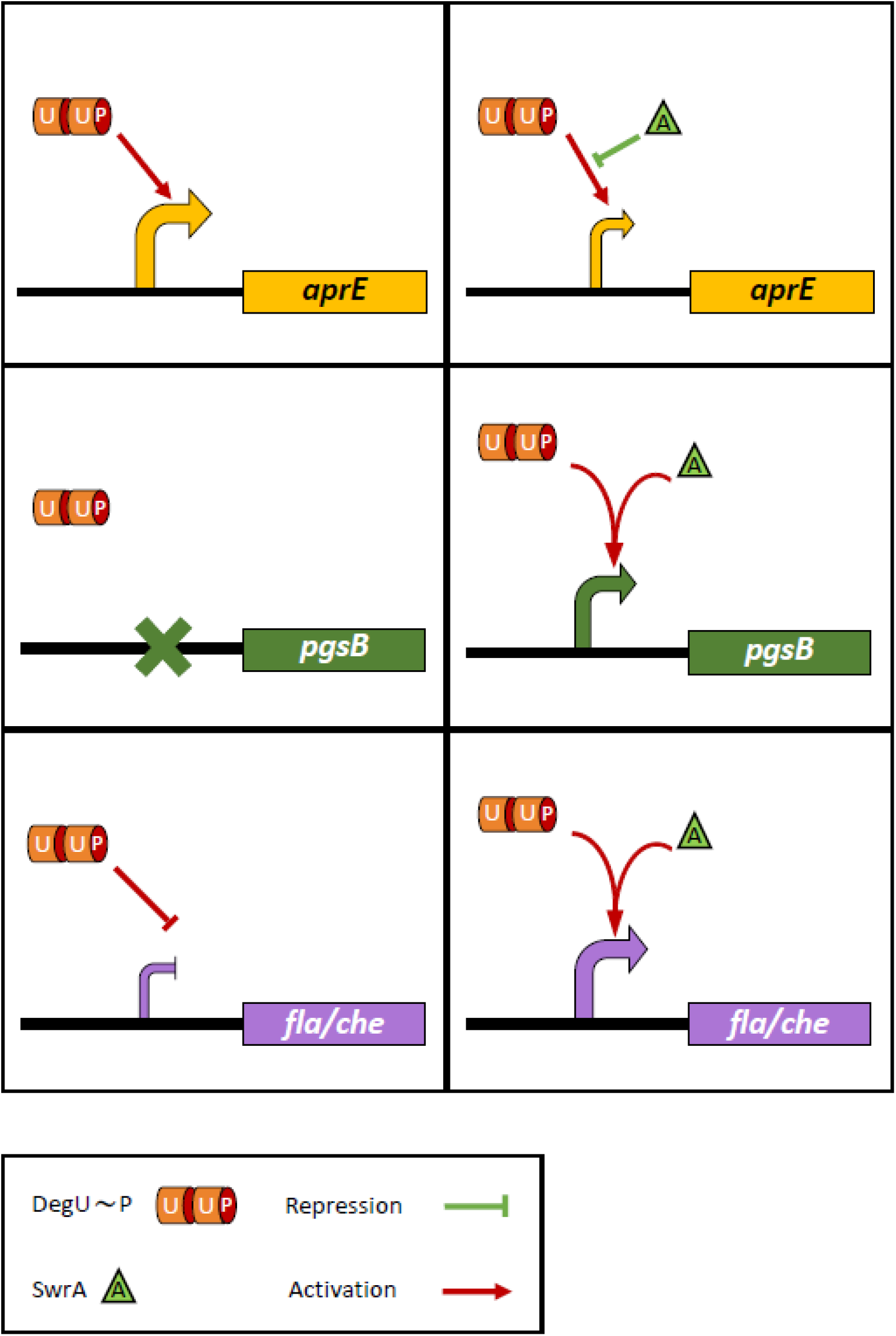
SwrA effects on DegS/U regulated genes. The schematic representations above summarizes the data collected on *aprE, pgs* and *fla/che* transcription on the global effects exerted DegU~P alone (left) and with SwrA (right). The size of the curved arrows in front of each gene is indicative of the efficiency of transcription. Symbols are described in the box at the bottom.

The results shown in this work do not confute literature data obtained in swrA^-^ domestic *B. subtilis* strains (168, JH642, and others) (18). The non-transformability of *degU*^Hy^ as well as *degS*^Hy^ swrA^-^ strains is indeed validated in our experimental settings (Fig. 1). Rather, a piece of literature data appears to support our results. In 1991, Hahn and Dubnau, analyzing the impact of *degU32*^Hy^ and *degS200*^Hy^ alleles on PsrfA expression, could not interpret the fact that, differently from DegU^Hy^, DegS^Hy^ did not repress *srfA* transcription (34). It is tempting to imagine that a high percentage of *swrA*^+^ revertant cells arose in the DegS^Hy^ strain used in the experiment, due to the high frequency of phase variation events (10^-4^) (17), and in those revertants SwrA was able to mitigate -or supress-the negative effect of DegU~P, turning it into a less negative -or positive-signal.

Presently, the main target of DegU~P in competence has not been clearly identified, because of the coexistence of at least two possible target genes: P_comk_ (8, 27, 28) and P_srfA_ (28, 34). A negative effect of the *degQ*^Hy^ allele has been evidenced on both promoters, in domestic and undomesticated strains (*swrA*^-^ and *swrA*^+^, respectively) (28). Since the *degQ*^Hy^ mutation increases DegU~P levels (5) without impacting on the DegU structure, its interaction with SwrA is preserved. The way in which the effects of SwrA and DegQ are balanced needs to be further analyzed in well-defined genetic backgrounds.

A second fundamental result that emerges from this work is that the DegU32^Hy^ mutant protein does not behave as the phosphorylated wild-type DegU protein. A proxy for DegU~P is represented by the *degS200*^Hy^ mutation, which produces high levels of DegU~P without directly modifying the structure of the transcriptional activator DegU. Moreover, from the lack of activation of the *pgs* promoter in undomesticated strains, which are naturally *swrA*^+^ *degQ*^Hy^ (data not shown), it can be hypothesized that the level of DegU phosphorylation attained in *degQ*^Hy^ cells is lower compared to that gained with the *degS200*^Hy^ mutation.

From P_pgs_ analyzes it can be hypothesized that DegU32^Hy^ is able to bind directly to DNA, even before activation of the signalling pathway that would lead to its phosphorylation, i.e., in the non-phosphorylated form (see Fig. 3A). This notion is not novel: Stanely-Wall and collaborators showed that γ-PGA production in a *degU32*^Hy^ background also occurs in a *degS* null mutant (35). Also *in vitro,* Mordini et al. (2013) showed that DegU32^Hy^ binds to DNA independently from the presence of its cognate kinase. Moreover, the interaction of SwrA (physical or genetical) with the mutant DegU32^Hy^ protein is compromised. Either it does not occur at all, as it appears by the lack of differences between the phenotypes of *degU32*^Hy^ *swrA*^+^ and *swrA*^-^ strains in competence, *aprE* expression and motility (Figs. 1 & 2 and ref. 16), or it markedly differs from the interaction with DegU~P produced by *degS200*^Hy^, as it appears from the differential activation profile of P_pgs_ in the two mutants. In any case, the physiological role of DegU~P in *B. subtilis* should be approached using a DegS200^Hy^ mutant. This also suggests that our current view of the impact of the DegS/U on *B. subtilis* physiology gained through the use of *degU32*^Hy^ mutants might require some revamping, as it happened for motility.

## MATERIALS AND METHODS

### Strain construction

All strains used in this study are listed in Table 1.

PB5630, corresponding to strain DK1042 obtained by D. Kearns and co-workers (29) by introducing the *com/*^Q12L^ mutation in the resident plasmid of the undomesticated NCIB3610, was transformed with pLoxSpec/degSU(Hy) and pLoxSpec/degS200 (24). PB5814 and PB5815, respectively, were obtained after selection for spectinomycin resistance (60 μg/ml).

**Table 1.**
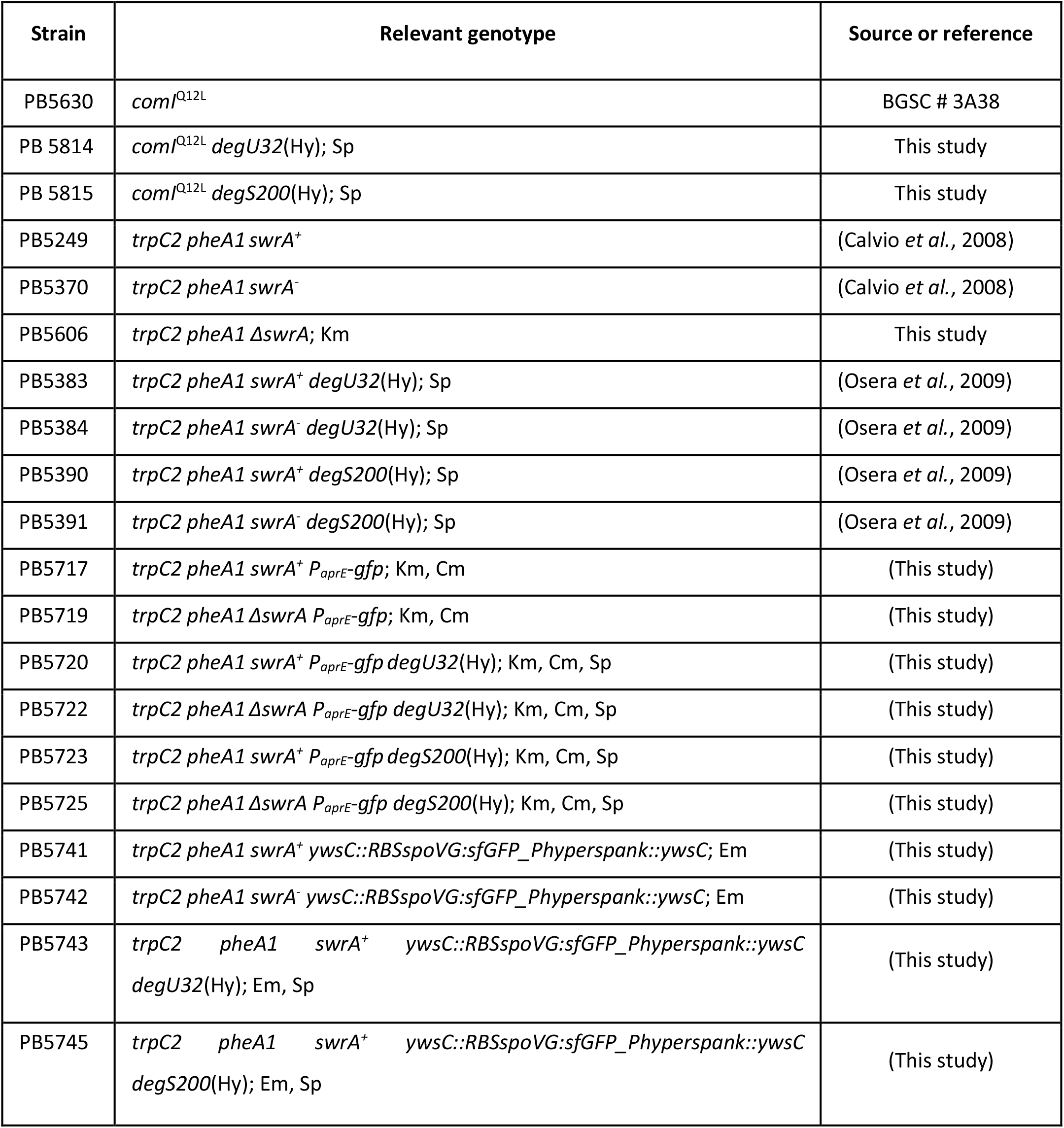

**Table 2.**

|  |  | Effect of DegU~P [*degS*(Hy)] |  |  |
| --- | --- | --- | --- | --- |
| Target | *degS*/*U*<sup>wt</sup>*swrA*<sup>-</sup> | *swrA*<sup>-</sup> | *swrA*<sup>+</sup> |  |
| P<sub>*fla*/*che*</sub> | motile | NEGATIVE (non-motile) | POSITIVE (hyperflagellation) | Mordini et al,<br>2013 |
| P<sub>*aprE*</sub> | no expression | POSITIVE (production) | PARTIALLY NEGATIVE (-38%) | Fig. 2 |
| P<sub>*pgs*</sub> | no expression | NONE | POSITIVE (mucoid colonies) | Fig. 3 |
| Competence | competent | NEGATIVE (non-competent) | PARTIALLY POSITIVE (+37%) | Fig. 1 |

The clean deletion of the *swrA* gene was obtained by transforming PB5249 with pCCΔswrA, a non-replicative plasmid that, completely removing the *swrA* ORF, inserts a kanamycin resistance gene upstream of the *swrA* promoter to control the expression of the downstream *minJ* gene. pCCΔswrA plasmid was obtained through the following steps: a PCR fragment comprising the region upstream the *swrA* gene, containing all the regulatory elements, was amplified from PB5249 genomic DNA with primers UPPromA/E (*Eco*RI)5’-cc<u>gaattc</u>tttgtgcttaaagagattatggatc-3’ and CC_A_rev (*Xho*I) 5’-aacg<u>ctcgag</u>ttgtgaacccccattttctttatacagataagcac-3’; the initial part of the following ORF, *minJ,* was amplified from the same source with primers CC_B_for (*Xho*I) 5’-acc<u>gctcgagg</u>tgtctgttcaatggggaattgaactgttaaaaagc-3’ and CC_C_rev (*Sma*I) 5’-tc<u>ccccgggg</u>tttgccagctgctgtccgatcg-3’. The two products were digested with *Xho*I (restriction sites underlined) and ligated. The 934 bp resulting product was inserted between the *Eco*RI and *Sma*I sites of the pJM114-derived pCC1 (21). The plasmid pCCΔswrA, verified by multiple restriction digestion and by sequencing of the relevant portions.

The plasmid pCCΔswrA was used to transform PB5249. PB5606 was obtained by selecting one clone for kanamycin (2 μg/ml) resistance; deletion of the coding sequence of *swrA* and the integrity of its promoter and *minJ* were verified by PCR and DNA sequencing.

The P_aprE_-*gfp* strains were obtained by *in^-^locus* integration of the pGFP-aprE plasmid (a generous gift from Prof. J.W. Veening, ref. 32) into the chromosome of *swrA*^+^ and *ΔswrA* isogenic strains, respectively PB5393 (21) and PB5606 (described above), both carrying a kanamycin resistance gene upstream of the *swrA* promoter region. The resulting strains were named PB5717 and PB5719, respectively (Table 1). *degU32*(Hy) and *degS200*(Hy) alleles, were introduced in PB5717 and PB5719 by transformation with pLoxSpec/degSUHy) and pLoxSpec/degS200 (24) and selection for spectinomycin resistance (60 μg/ml). In the derived strains, PB5720, PB5722, PB5723 and PB5725 (Table 1), the single copy insertion of each construct was assessed.

The P_pgs_-SF*gfp* strains were obtained using a modified pMutin vector (pMATywsC). The construction of the plasmid occurred in multiple steps. First, in the pMutin-GFP vector (ECE149, obtained from the Bacillus genetic stock centre, http://www.bgsc.org/) the *gfp* gene was substituted by Gibson assembly with a super folder version of the GFP (SFgfp) amplified from pECE323 plasmid (Bacillus genetic stock centre) with primers RXeGFPda321-5’-ggctgcactagtgctcgaattcattatttataaagttcgtccataccgtg-3’ and FXeGFPda321-5’-tcggccggaaggagatatacatatgtcaaaaggagaagaactttttacag-3’ to give pMutinsfGFP. The 5’ portion of *ywsC* together with the Phyperspank promoter were inserted in the resulting pMutinsfGFP through a tripartite Gibson assembly. The Phyperspank promoter was amplified from plasmid Phyp.R0.sfGFP(sp).LacI_operon (36) using primers PHypFor 5’-agcttccaagaaagatatccctcggatacccttactctgttg-3’ and PHypRev 5’-ggctataatgagtaaccacatgtttgtcctccttattagttaatc-3’; the 5’ portion of *ywsC* (647 bp) was amplified from PB5249 chromosomal DNA using primers ywsCFor 5’-taactaataaggaggacaaacatgtggttactcattatagcctgtg-3’ and ywsCRev 5’-gtaaaaagttcttctccttttgacagagaagcgttatcagggaatac-3’. In the plasmid obtained, pPhsywsCsfGFP, the spoVG RBS and initial codons were translationally fused to the sfGFP using the partially overlapping oligos oligoFORSpoVG 5’-ccctgataacgcttctctggaattcccgggatccccagcttgttgatacactaatgcttttatatagggaaaaggtggtgaactactatgTCAAAAGGAG-3’ and oligoREVSpoVG 5’-CTCCTTTTGAcatagtagttcaccaccttttccctatataaaagcattagtgtatcaacaagctggggatcccgggaattccagagaagcgttatcaggg-3’, derived from the pJM116 vector (37) The final construct was verified by sequencing and saved as pMATywsC. This plasmid was used to transform PB5249 (*swrA*^+^) and PB5370 (*swrA*^-^), using erythromycin resistance (5 μg/ml) for selection, resulting in PB5741 and PB5742 strains, respectively. *degU32*(Hy) and *degS200*(Hy) alleles were introduced in PB5741 by transformation with pLoxSpec/degSU(Hy) and pLoxSpec/degS200 (24) by spectinomycin resistance (60 μg/ml) selection, giving rise to PB5743 and PB5745, respectively. The single copy insertion of each construct was assessed.

### Competence evaluation

Cells were inoculated in LM (LB supplemented with MgSO_4_, 9μM; tryptophan, 50 μg/mL; phenylalanine, 50 μg/mL) at OD_600_=0.2 and grown at 37°C shaking. At OD_600_=1, cells were diluted 1:20 in MD (K_2_HPO_4_, 9.8 mg/ml; KH_2_PO_4_, 5.52 mg/ml; Na_3_Citrate·5H_2_O, 0.92 mg/ml; glucose, 20 mg/ml; tryptophan, 50 μg/ml; phenylalanine, 50 μg/ml; ferric ammonium citrate, 11 μg/ml; K-aspartate, 2.5 mg/ml; MgSO_4_, 0.36 mg/ml) and grown at 37°C until stationary phase (T_0_). About 200 ng of chloramphenicol (Cm)-selectable *B. subtilis* chromosomal DNA was added to 0.5 ml cells which were further incubated for 1.5 h at 37°C with shaking. Transformants were isolated on 5 mg/ml chloramphenicol on several selective plates. Resistant colonies were counted and related to cell density at T_0_ to calculate the transformation efficiency, taking into account each dilution step before plating. Data shown in Fig. 1 represent the average of three independent experiments.

### Gene expression evaluation by flow cytometry

For the analysis of *P*_aprE_ activity, cells were inoculated in Shaeffer’s sporulation medium (38) at 0.2 OD_600_ and grown at 37°C under continuous shaking for 20 h. Aliquots were collected every 60’ for OD_600_ readings; at the transition point (5h), 5 and 15 h later, 10% glycerol (final concentration) was added to culture aliquots for storage at −20°C.

For the analysis of *P*pgs activity, cells were inoculated in E-medium (39) at OD_600_=0.1 and grown at 37°C under continuous shaking for 48h. Aliquots were collected at 2-h intervals for OD_600_ readings and direct cytofluorimetric analyses. Before analyses, fresh and/or frozen samples were centrifuged for 5 minutes at 14000 xg; cell pellets were re-suspended in D-PBS for flow cytometry (Gibco).

Samples were acquired on an Amnis® ImageStream®X Mk II Imaging Flow Cytometer using the INSPIRE software with the following set up: Channel 02 (GFP fluorescence), Channel 06 (scattering channel); the Brightfield was visualized on Channel 01 and on Channel 05, depending on the GFP expression level, to avoid interference from Channel 02. The 488 nm laser was used at either 50 mW or 100 mW power, according to the GFP expression level, in order to avoid sensor saturation. The flow rate was set to low speed/high sensitivity and images were taken at 60X magnification. For each sample at least 10000 events were acquired.

All data were analyzed using the IDEAS software (version 6.2). In-focus events were gated in a histogram displaying the Gradient RMS_M01_Ch01 on the x-axis. A plot of the Area versus Aspect Ratio Intensity in the Brightfield channel was used to exclude doublets from the analysis. A plot of the Area versus Intensity in the Scattering channel was used to exclude events characterized by high scatter such as beads. To avoid any bias due to cell size in evaluating fluorescence intensity, the GFP level of each cell was calculated through the Median Pixel feature on the fluorescence channel. The threshold value to distinguish the ON population was set at the maximum autofluorescence of a non-fluorescent population used as negative control (OFF). Data presented in Figs. 2 and 3 represent the average of three independent experiments.

## ACKNOWLEDGEMENTS

This research was supported by the Italian Ministry of Education, University and Research (MIUR): Dipartimenti di Eccellenza Program (2018–2022) - Dept. of Biology and Biotechnology “L. Spallanzani”, University of Pavia. Authors are grateful to Prof. J. W. Veening for the P_aprE_ plasmid. The contribution of Erlinda Rama, Matteo Cavaletti, Jessica Bollini, Laura Nucci is acknowledged. We also thank Dr A. Azzalin, from the departmental imaging facility, and Dr A. Serra, from Luminex Corporation, for support with the Amnis ImageStream data acquisition and analyses.

## SUPPLEMENTARY MATERIALS

**Fig. S1.**
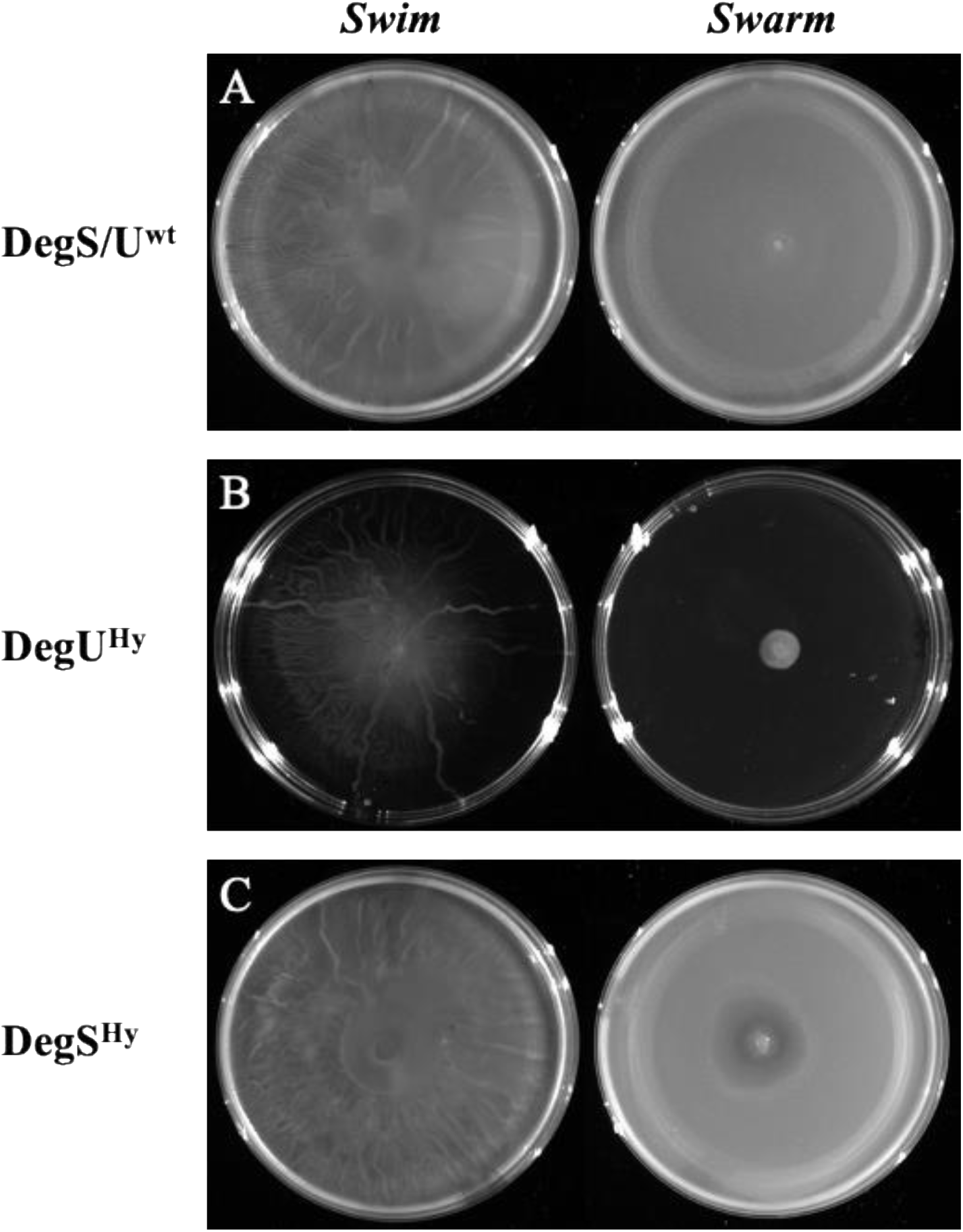
Motility of undomesticated DegU/S^Hy^ strains. Swimming (on the left of each panel) and swarming (on the right) motility performances of undomesticated strains (*swrA*^+^). A): PB5630 *degS/U*^wt^; B): PB5814 *degU*^Hy^; C) PB5815 *degS*^Hy^. The genotype of each strain is also indicated on the right. Strains are listed in Table 1. Motility plates were prepared as described in Mordini *et al.,* 2013, except for the addition of surfactin, that was omitted in the swarming plates.

**Fig. S2.**
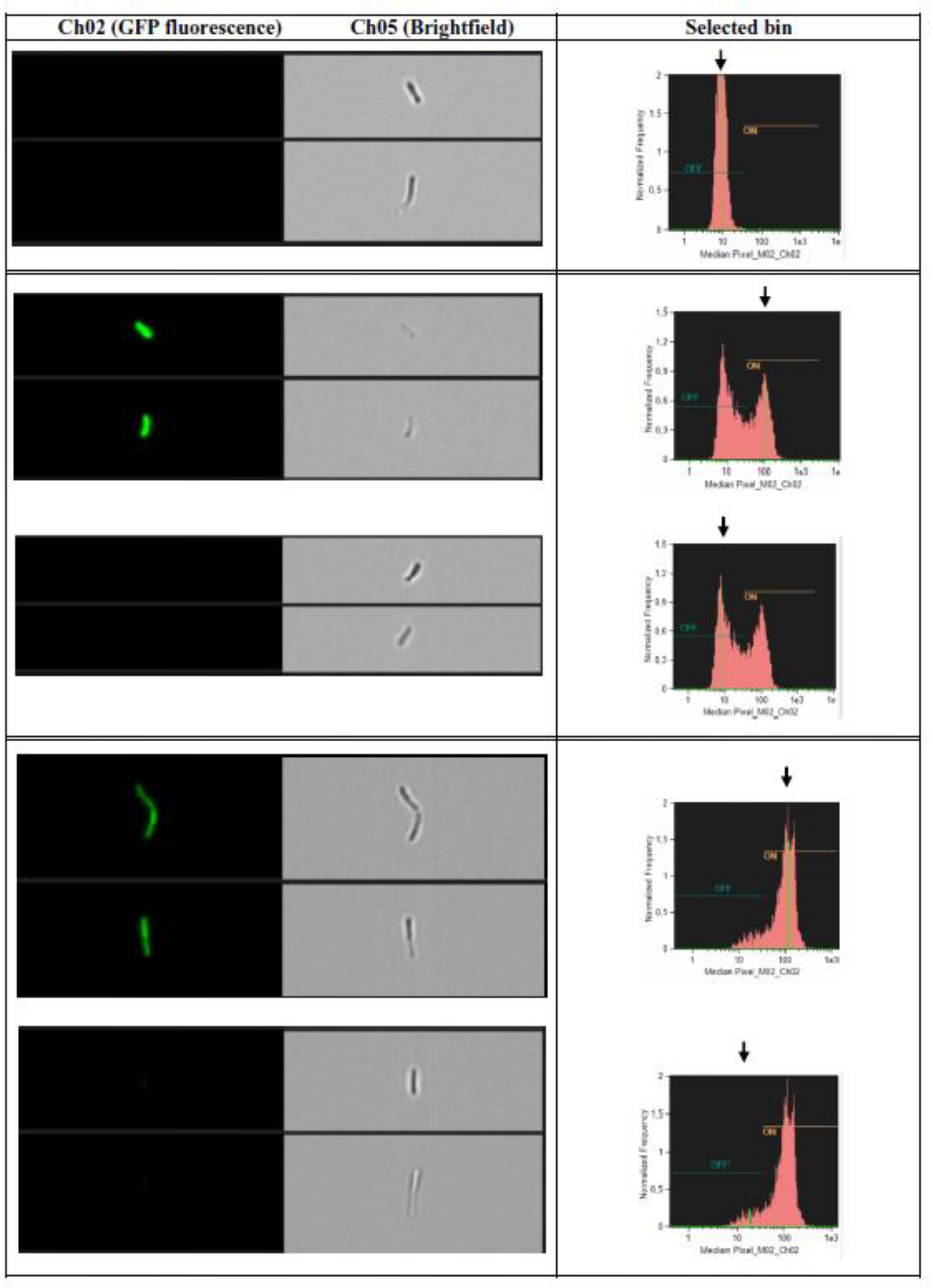
Cell images collected during Flow Cytometry. Representative images of P_aprE_-GFP containing cells collected from the different intensity bins which are pointed by an arrow on the graphs in the right panels. On the left, identical images as acquired in the fluorescence channel (Ch02) and in the brightfield channel (Ch05).

## Notes

### Competing Interest Statement

The authors have declared no competing interest.

## REFERENCES

1. Mitrophanov AY, Groisman EA. 2008. Signal integration in bacterial two-component regulatory systems. Genes & development 22:2601–2611.

2. Msadek T, Kunst F, Henner D, Klier A, Rapoport G, Dedonder R. 1990. Signal transduction pathway controlling synthesis of a class of degradative enzymes in Bacillus subtilis: expression of the regulatory genes and analysis of mutations in degS and degU. J Bacteriol 172:824–34.

3. Ogura M, Yamaguchi H, Yoshida Ki, Fujita Y, Tanaka T. 2001. DNA microarray analysis of Bacillus subtilis DegU, ComA and PhoP regulons: an approach to comprehensive analysis of B.subtilis two- component regulatory systems. Nucleic Acids Res 29:3804–13.

4. Mäder U, Antelmann H, Buder T, Dahl M, Hecker M, Homuth G. 2002. Bacillus subtilis functional genomics: genome-wide analysis of the DegS-DegU regulon by transcriptomics and proteomics. Mol Genet Genomics 268:455–67.

5. Kobayashi K. 2007. Gradual activation of the response regulator DegU controls serial expression of genes for flagellum formation and biofilm formation in Bacillus subtilis. Molecular Microbiology 66:395–409.

6. Kunst F, Pascal M, Lepesant-Kejzlarova J, Lepesant J, Billault A, Dedonder R. 1974. Pleiotropic mutations affecting sporulation conditions and the syntheses of extracellular enzymes in Bacillus subtilis 168. Biochimie 56:1481–9.

7. Dahl M, Msadek T, Kunst F, Rapoport G. 1991. Mutational analysis of the Bacillus subtilis DegU regulator and its phosphorylation by the DegS protein kinase. J Bacteriol 173:2539–47.

8. Dahl M, Msadek T, Kunst F, Rapoport G. 1992. The phosphorylation state of the DegU response regulator acts as a molecular switch allowing either degradative enzyme synthesis or expression of genetic competence in Bacillus subtilis. J Biol Chem 267:14509–14.

9. van Sinderen D, Venema G. 1994. comK acts as an autoregulatory control switch in the signal transduction route to competence in Bacillus subtilis. J Bacteriol 176:5762–70.

10. Amati G, Bisicchia P, Galizzi A. 2004. DegU-P represses expression of the motility fla-che operon in Bacillus subtilis. J Bacteriol 186:6003–14.

11. Mukai K, Kawata-Mukai M, Tanaka T. 1992. Stabilization of phosphorylated Bacillus subtilis DegU by DegR. J Bacteriol 174:7954–62.

12. Yang M, Ferrari E, Chen E, Henner D. 1986. Identification of the pleiotropic sacQ gene of Bacillus subtilis. J Bacteriol 166:113–9.

13. McLoon AL, Guttenplan SB, Kearns DB, Kolter R, Losick R. 2011. Tracing the domestication of a biofilm-forming bacterium. J Bacteriol 193:2027–34.

14. Henner DJ, Yang M, Ferrari E. 1988. Localization of Bacillus subtilis sacU(Hy) mutations to two linked genes with similarities to the conserved procaryotic family of two-component signalling systems. J Bacteriol 170:5102–9.

15. Tanaka T, Kawata M, Mukai K. 1991. Altered phosphorylation of Bacillus subtilis DegU caused by single amino acid changes in DegS. J Bacteriol 173:5507–15.

16. Mordini S, Osera C, Marini S, Scavone F, Bellazzi R, Galizzi A, Calvio C. 2013. The role of SwrA, DegU and P(D3) in fla/che expression in B. subtilis. PLoS One 8:e85065.

17. Kearns DB, Chu F, Rudner R, Losick R. 2004. Genes governing swarming in Bacillus subtilis and evidence for a phase variation mechanism controlling surface motility. Mol Microbiol 52:357–69.

18. Calvio C, Celandroni F, Ghelardi E, Amati G, Salvetti S, Ceciliani F, Galizzi A, Senesi S. 2005. Swarming differentiation and swimming motility in Bacillus subtilis are controlled by swrA, a newly identified dicistronic operon. J Bacteriol 187:5356–5366.

19. Zeigler DR, Prágai Z, Rodriguez S, Chevreux B, Muffler A, Albert T, Bai R, Wyss M, Perkins JB. 2008. The origins of 168, W23, and other Bacillus subtilis legacy strains. J Bacteriol 190:6983–95.

20. Kearns DB, Losick R. 2005. Cell population heterogeneity during growth of Bacillus subtilis. Genes Dev 19:3083–94.

21. Calvio C, Osera C, Amati G, Galizzi A. 2008. Autoregulation of swrAA and motility in Bacillus subtilis. J Bacteriol 190:5720–5728.

22. Urushibata Y, Tokuyama S, Tahara Y. 2002. Difference in transcription levels of cap genes for gamma-polyglutamic acid production between Bacillus subtilis IFO 16449 and Marburg 168. J Biosci Bioeng 93:252–4.

23. Stanley N, Lazazzera B. 2005. Defining the genetic differences between wild and domestic strains of Bacillus subtilis that affect poly-gamma-dl-glutamic acid production and biofilm formation. Mol Microbiol 57:1143–58.

24. Osera C, Amati G, Calvio C, Galizzi A. 2009. SwrAA activates poly-gamma-glutamate synthesis in addition to swarming in Bacillus subtilis. Microbiology-Sgm 155:2282–2287.

25. Ogura M, Tsukahara K. 2012. SwrA regulates assembly of Bacillus subtilis DegU via its interaction with N-terminal domain of DegU. J Biochem 151:643–655.

26. Verhamme D, Kiley T, Stanley-Wall N. 2007. DegU co-ordinates multicellular behaviour exhibited by Bacillus subtilis. Mol Microbiol 65:554–68.

27. Hamoen LW, Van Werkhoven AF, Venema G, Dubnau D. 2000. The pleiotropic response regulator DegU functions as a priming protein in competence development in Bacillus subtilis. Proc Natl Acad Sci U S A 97:9246–51.

28. Miras M, Dubnau D. 2016. A DegU-P and DegQ-Dependent Regulatory Pathway for the K-state in Bacillus subtilis. Frontiers in Microbiology 7:1868.

29. Konkol MA, Blair KM, Kearns DB. 2013. Plasmid-Encoded ComI Inhibits Competence in the Ancestral 3610 Strain of Bacillus subtilis. J Bacteriol 195:4085–4093.

30. Srivatsan A, Han Y, Peng J, Tehranchi AK, Gibbs R, Wang JD, Chen R. 2008. High-precision, whole-genome sequencing of laboratory strains facilitates genetic studies. PLoS Genet 4:e1000139.

31. Henner DJ, Ferrari E, Perego M, Hoch JA. 1988. Location of the targets of the hpr-97, sacU32(Hy), and sacQ36(Hy) mutations in upstream regions of the subtilisin promoter. J Bacteriol 170:296–300.

32. Veening J, Igoshin O, Eijlander R, Nijland R, Hamoen L, Kuipers O. 2008. Transient heterogeneity in extracellular protease production by Bacillus subtilis. Mol Syst Biol 4:184.

33. Veening J, Hamoen L, Kuipers O. 2005. Phosphatases modulate the bistable sporulation gene expression pattern in Bacillus subtilis. Mol Microbiol 56:1481–94.

34. Hahn J, Dubnau D. 1991. Growth stage signal transduction and the requirements for srfA induction in development of competence. J Bacteriol 173:7275–7282.

35. Cairns LS, Marlow VL, Bissett E, Ostrowski A, Stanley-Wall NR. 2013. A mechanical signal transmitted by the flagellum controls signalling in Bacillus subtilis. Mol Microbiol 90:6–21.

36. Guiziou S, Sauveplane V, Chang HJ, Clerté C, Declerck N, Jules M, Bonnet J. 2016. A part toolbox to tune genetic expression in Bacillus subtilis. Nucleic Acids Res 44:7495–508

37. Perego M. 1993. Integrational Vectors for Genetic Manipulation in Bacillus subtilis, p 615–624. In Bacillus subtilis and Other Gram-Positive Bacteria (eds A.L. Sonenshein, J.A. Hoch and R. Losick).

38. Harwood CR, Cutting SM. 1990. Molecular Biological Methods for Bacillus. John Wiley and Sons, Chichester.

39. Scoffone V, Dondi D, Biino G, Borghese G, Pasini D, Galizzi A, Calvio C. 2013. Knockout of pgdS and ggt genes improves gamma-PGA yield in B. subtilis. Biotechnol Bioeng 110:2006–12.

